# Engaging narratives evoke similar neural activity and lead to similar time perception

**DOI:** 10.1101/104778

**Authors:** Samantha Cohen, Simon Henin, Lucas C. Parra

## Abstract

It is said that we lose track of time - that “time flies” - when we are engrossed in a story. How does engagement with the story cause this distorted perception of time, and what are its neural correlates? People commit both time and attentional resources to an engaging stimulus. For narrative videos, attentional engagement can be represented as the level of similarity between the electroencephalographic responses of different viewers. Here we show that this measure of neural engagement predicted the duration of time that viewers were willing to commit to narrative videos. Contrary to popular wisdom, engagement did not distort the average perception of time duration. Rather, more similar brain responses resulted in a more uniform perception of time across viewers. These findings suggest that by capturing the attention of an audience, narrative videos bring both neural processing and the subjective perception of time into synchrony.

## Introduction

Most people are familiar with the experience of becoming completely captivated by a book or movie ^1^. This phenomenon has been described as “engagement”, “transportation”, “absorption”, or “flow” ^2–4^. Some argue that when we are fully absorbed in a narrative, there is a loss of conscious awareness of the external environment and the passage of time ^2,5,6^. There are contradicting theories regarding whether engaging stimuli will elongate or condense perceived time. For instance, if a stimulus is enjoyable, it may evoke greater attention, thus creating a richer perceptual experience that seems longer in retrospect ^7,8^. On the other hand, when a stimulus commands attention, it may distract attention away from the passage of time, thus decreasing the perceived time elapsed ^9^. Further complicating matters, the nature of the temporal distortion is likely dependent on the emotional valence of the stimulus ^10^.

Here we will explore the relationship between attentional engagement with video narratives and the perceived passage of time on the scale of seconds. Previous research on the neural basis of time perception has examined durations that range from milliseconds to days^11^. For stimuli in the order of seconds, previous work mostly concerns controlled stimuli, such as constant tones or images ^12^. Given the importance of attention to time perception ^13^, it is possible that time perception may be substantially different during more realistic scenarios, such as engaging narratives ^14^. Attention is also known to modulate the similarity of electroencephalographic (EEG) evoked responses across viewers during narrative videos ^15^. In this paper, attentional engagement was assessed from the inter-subject correlation (ISC) of EEG responses ^16^. For an objective reference of stimulus engagement, we also define a behavioral measure of engagement that is based on the time that online viewers commit to watching the narrative videos. As we will show, this is an objective, value-based metric. Unlike previous measures ^5,17^, it is independent of subjective self-report biases and its assessment does not interrupt the processing of the stimulus. Engagement, whether measured neurally, or behaviorally, is thus defined by the commitment to devote a scarce resource to the stimulus. In this case, that resource is either attention or time, and, as predicted, these engagement measures are correlated. Next, we address whether moments of high engagement prolong or shorten the perception of time. Surprisingly, neither the behavioral measure of time commitment, nor the neural measure of attention coincided with the perceived passage of time. Instead, the similarity of brain responses predicted the uniformity of time estimates across viewers. This robust effect was reproduced across two cohorts of viewers. Thus, engagement does not appear to distort perceived time duration, but rather, engagement, inducing a more uniform neural processing of the stimulus, leads to a more uniform assessment of time.

## Results

We define engagement as the commitment to devote a scarce resource, such as attention or time, to a stimulus. Three experiments were performed to establish behavioral and neural measures of engagement, and to relate these assessments to time perception. In the first experiment, behavioral engagement was evaluated from the viewing behavior of large online audiences. In the second experiment, neural engagement was extracted from the EEG responses to the same videos for which behavioral engagement had been measured. In the third experiment, the perception of time was queried during short intervals within the videos.

### 1. Experimental assessment of engagement behavior

Time commitment can be calculated from the stimulus’s ability to retain viewers. For a large enough audience, this can be measured from viewership survival, S(t), defined as the fraction of the audience that “survives” until time, t, in the video (Figure 1A). The rate at which the audience shrinks is the risk of viewership loss, denoted here as *λ*(*t*) (Equation 3 in Methods Section 3, and Supplementary Fig. S1). When the stimulus evokes a high level of engagement, the risk of losing viewers is low. Conversely, when the audience is not engaged, the risk is high. Instantaneous behavioral engagement is formally defined as the inverse of the risk of losing viewers:

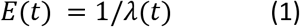

**Figure 1:**
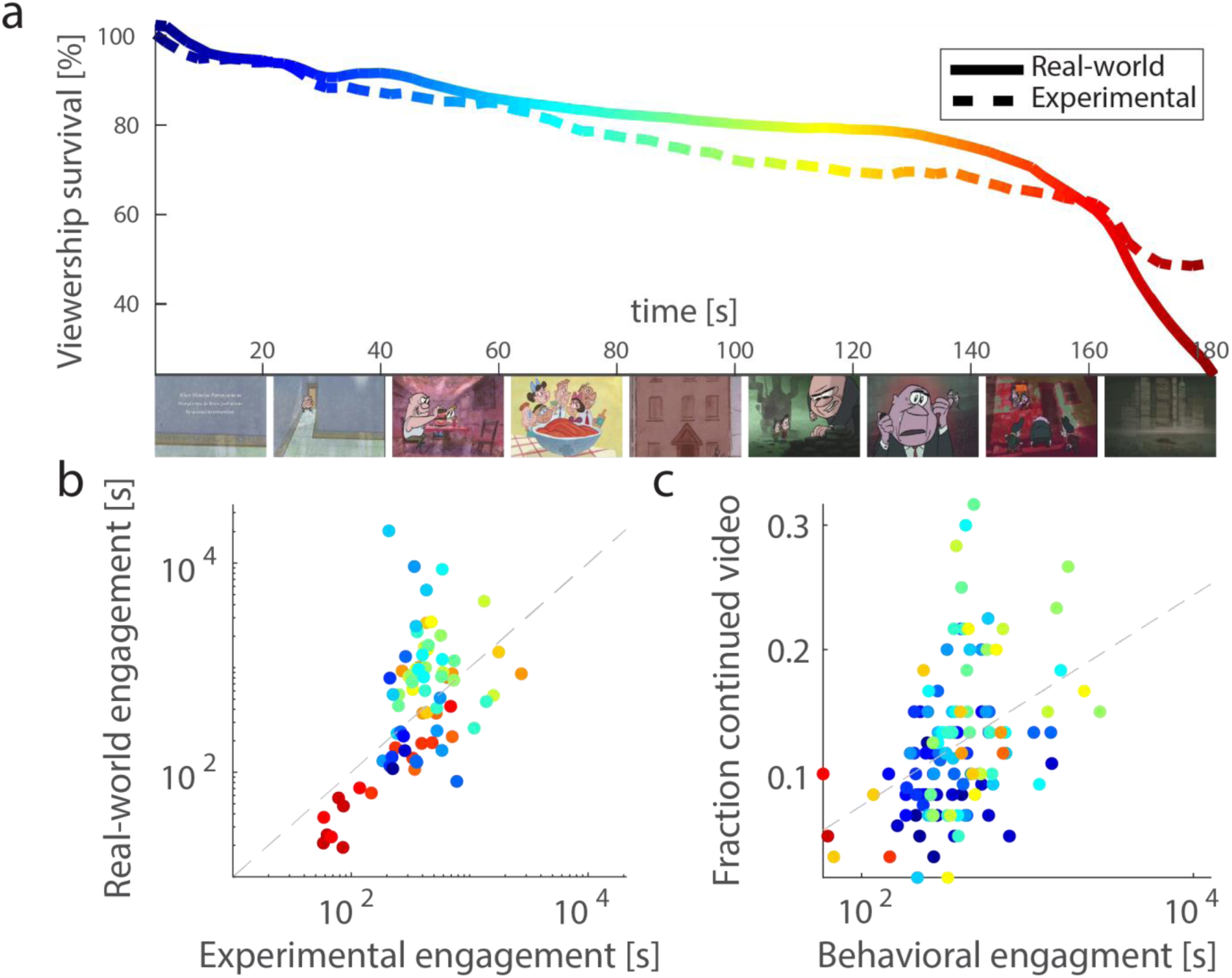
Behavioral engagement in “experimental” cohort mimics “real-world” behavior. A: Viewership survival, shown for an example video, is measured as the fraction of the initial number of viewers who are retained over time. “Real-world” data (darker line) was provided for this study by its content owner, StoryCorps, and represents the viewership (239,511 views) accumulated over several years for this video. “Experimental” survival (lighter line) was collected in approximately one hour from 1000 viewers recruited online via Amazon’s Mechanical Turk (MTurk) platform. Still images from “Sundays at Rocco’s,” a StoryCorps animated short directed by the Rauch Brothers and produced by Lizzie Jacobs and Mike Rauch, reproduced here with permission from StoryCorps. B: Variation in engagement across time correlates between real-world and experimental data (r=0.60, p=2e-7, N=77). Engagement was estimated using a time interval of ∆t=12s for the five videos that were common to both conditions. Dashed line indicates points with equal engagement in both data-sets. C: Experimental engagement correlated with the likelihood that a separate cohort of viewers voluntarily continued to watch the videos when given the option to stop. Dashed line represents the regression line. In both B and C each point represents a time interval, color represents the corresponding time point in the video from A, and engagement is displayed on a log-seconds scale.

In the present context, where time is the scarce resource being allocated, E(t) is equivalent to the additional time, measured in seconds, that the average viewer is willing to invest in the stimulus (for more detail on interpretation, see Methods Section 3 and Discussion).

Raw viewership survival data (Figure 1A) was collected for two cohorts of viewers. One cohort voluntarily watched the video stimuli online (real-world, 5 videos, approximately 2 million viewers over 2.6 +/- 0.8 years, data shared by StoryCorps). The second cohort consisted of a group of subjects who were directed to the videos as part of an experimental paradigm (experimental, 10 videos, N=1000, collected over approximately 1 hour on Amazon’s Mechanical Turk platform, MTurk; see Methods). All video stimuli were animated renditions of biographical narratives (161 ± 44 s, mean ± standard deviation). Five of the videos were viewed by both cohorts. In the real-world, viewers commit time to watch videos despite real-world commitments and time pressures. In contrast, experimental viewers were given an artificial time pressure of 15 minutes to access 27 minutes of video. For the videos common to both groups, real-world viewers were found to be significantly more engaged than those recruited experimentally (t(4)=3.2, p=0.03, paired t-test), Supplementary Fig. S2).

Despite this disparity in overall engagement, the two groups exhibited a similar modulation in engagement over the course of the videos. This relationship was present regardless of the time scale (∆t) at which engagement was evaluated (in Equation 3). The correlation between the experimental and real-world datasets was stable for time intervals, ∆t, ranging from 4 to 21 seconds (r = 0.57 +/- 0.06). Here, and in all subsequent analyses, time and engagement were measured on a logarithmic scale following convention in time perception ^18^ and survival analysis literature ^19^. Figure 1B displays this relationship for a time interval of ∆t = 12 s. While in the real-world viewership drops by approximately 9,000 viewers in an interval of 12 s, in the experimental cohort only 7 viewers are lost. Therefore, the experimental assessment of engagement is a noisier metric (see Methods, Section 3 and compare Supplementary Fig. 1). To validate the interpretation of this measure of engagement as a time commitment, we assessed the willingness to continue to watch the videos in a separate experiment (Results Section 3). Since this additional cohort was being compensated for performing a different task, the decision to continue to watch a video represents both a time and a financial sacrifice. Engagement measured experimentally was correlated with the fraction of viewers that voluntarily elected to continue to watch the videos after completing their time estimation task (r = 0.40, p = 5e-6, N = 121, Figure 1C). These independent validations indicate that the experimentally derived measure of engagement is a good data-set with which to evaluate a neural measure of engagement.

### 2. Neural engagement predicts behavioral engagement

We previously proposed that the similarity of electroencephalographic (EEG) evoked responses across viewers may be a neural marker of engagement ^16^. The similarity of EEG activity across subjects can be measured as the inter-subject correlation (ISC) of stimulus evoked responses ^16^. Since ISC is sensitive to attentional state ^15^, we predicted that there would be a relationship between ISC and the behavioral measure of engagement. To calculate ISC, EEG was recorded from 20 individuals who watched the same 10 videos from above. Components of maximal inter-subject correlation were then extracted from the EEG (see Methods). These components, C1-C3, capture sources of the evoked neural responses that are correlated in time across the entire sample of viewers (corresponding spatial distributions shown in Figure 2A). As such, each component potentially captures a different aspect of neural processing (e.g. visual, auditory, or supramodal processing, ^20^). The ISC of each component can be resolved in short time intervals during the stimulus ^16,21^, and time-resolved ISC was used to predict time-varying behavioral engagement.

**Figure 2:**
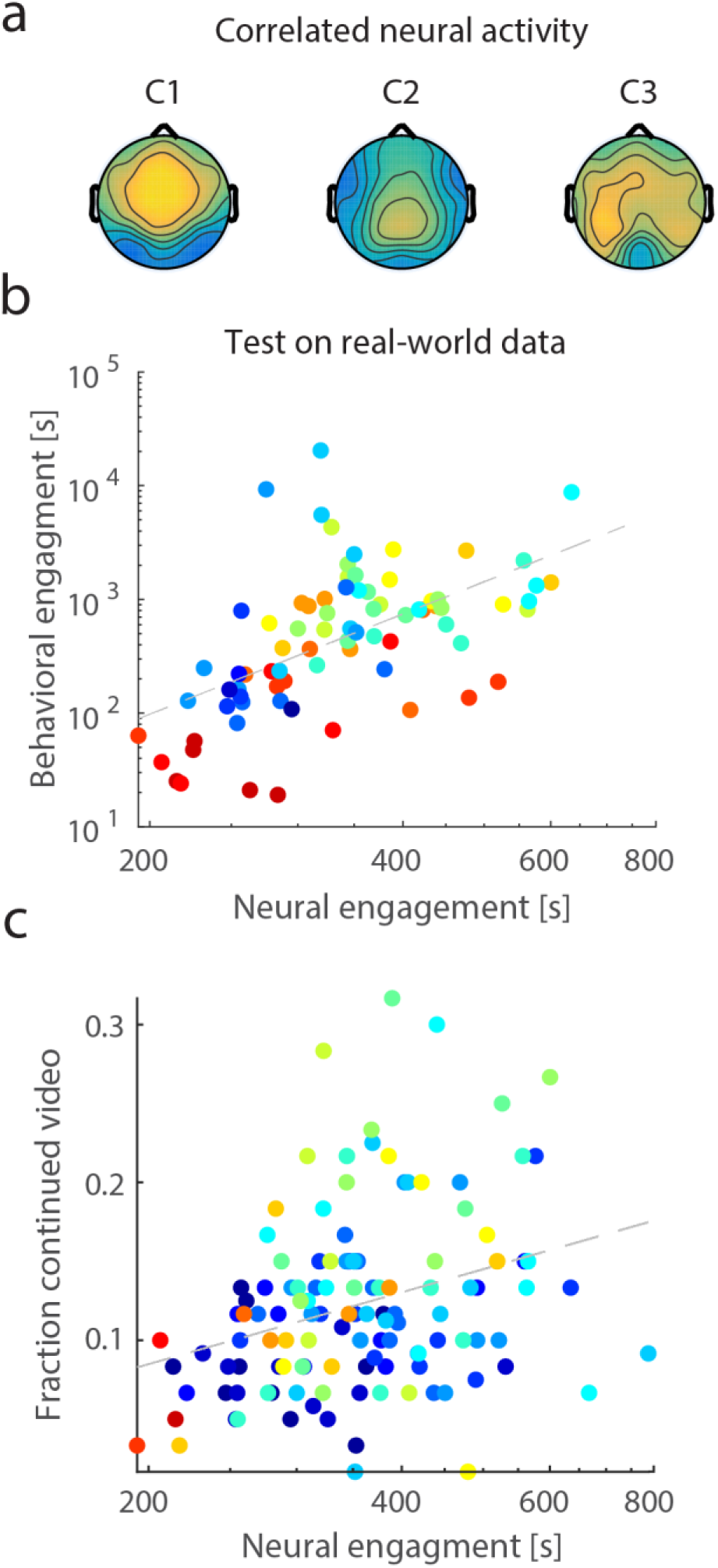
Neural Engagement predicts Behavioral Engagement. A: Spatial distribution of the three EEG components with maximal inter-subject correlation (ISC; C1 - C3). Color indicates positive (yellow) or negative (blue) correlation between the source of the neural responses and each sensor on the scalp. Component C2, center, contributes most to the relationship between neural and behavioral engagement (see main text). B: Neural engagement *Ê*(*t*); dashed line) predicts real-world behavioral engagement, *E*(*t*) using the model developed on the experimental behavioral engagement data (r = 0.56, p = 8e-8, N=78, for the 5 testing videos, ∆t=12s). C: Neural engagement correlated with the fraction of a separate cohort of viewers that decided to continue to watch the videos when given the option to stop prematurely. Dashed line represents the regression line (r=0.31, p=6e-4, N=122 intervals from all 10 videos). Each point represents a different time interval in each video colored according to time as in Figure 1. Both engagement measures are displayed on a log-seconds scale.

A regression model was first fit to the experimental behavioral engagement data (see Methods Section 7, Equation 4). This model’s predictive ability was then tested on the real-world data. Goodness of fit was assessed for different time intervals, ∆t, over which both behavioral engagement and ISC were calculated (Supplementary Fig. S3). ∆t=12s was selected as a good compromise between performance and number of samples (i.e. this ∆t had the smallest p-value, p=2e-6 with, R=0.4, N=128). The predictor of engagement behavior, Ê(*t*), which we refer to as “neural engagement” can be written as a product of baseline engagement, *E*_0_, with a time varying neural factor, *γ*(*t*):

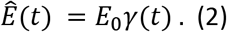

The baseline level of engagement, *E*_0_, was estimated to be 212 s, and the overall estimated engagement, averaged in time for all videos, was 307 s. Thus, for the experimental behavioral data, approximately 30% of the commitment to watch the stimuli could be accounted for by the temporal variation of the neural predictor variable *γ*(*t*) (log-sum of ISC in the largest three components; Equation 4). Behavioral engagement was mostly explained by the ISC of the second component (C2 in Figure 2A), which scales baseline engagement by a factor of 1.5 +/- 0.3 (mean and std of *γ*_2_(*t*) in Equation 4, Methods). Components C1 and C3 of the ISC contributed relatively less (1.0 +/- 0.005 and 1.1 +/- 0.1, mean and std of *γ*_1_(*t*)and *γ*_3_(*t*) respectively in Equation 4).

To test how well this neural engagement model predicts unseen data, we compared it to the real-world behavioral engagement data and found a significant correlation (r = 0.56, p = 8e-8, N=78 intervals from 5 videos, Figure 2B). In fact, this correlation was equally strong when training the regression coefficient with the experimental data from the five videos that were not part of the real-world behavioral data (r = 0.58, p = 2e-8, N=78). Thus, the predictive neural engagement model not only generalizes to unseen data, but it also generalizes across different stimuli. As with the behavioral engagement measures, the neural engagement measure, assessed across all videos, also correlated with the voluntary election to continue to watch the videos during the time estimation task (r=0.31, p=6e-4, N=122 intervals from all 10 videos, Figure 2C). The consistency between the neural and behavioral measures confirms the hypothesis that the similarity of brain responses captures attentional engagement.

### 3. Relationship between engagement and time perception

After establishing the validity of both the behavioral and neural measures of engagement, the relationship between stimulus engagement and time perception was assessed. An additional cohort of viewers (recruited from MTurk) provided subjective estimates for the durations of brief periods of time during the videos. These segments corresponded to those for which behavioral and neural engagement were assessed (∆t=12s, Figures 1 and 2). Each video was shown until an interval of interest, indicated by a visual cue, and after the time interval had elapsed, subjects were asked to report the perceived duration within a range of 8s to 16s (“Restricted range”, see Methods for implementation details).

Consistent with existing literature ^13,22^, the duration of the 12 s intervals was underestimated (11.3 +/- 0.03 s), and time intervals later in the videos were perceived as lasting longer than earlier intervals (r = 0.57, p = 2e-12, Supplementary Fig. S4). However, despite prevalent theories that stimulus engagement induces time distortion ^17^, there was no correlation between the mean estimates of time duration and engagement, measured either behaviorally or neurally (p > 0.3). Interestingly, however, both neural and behavioral engagement correlate with the *variability* of time estimates across viewers (Figure 3, Restricted range, for neural engagement: r = - 0.27, p=2e-3, N = 129, and for real-world behavioral engagement: r = - 0.25, p=0.03, N = 78 intervals from the 5 real-world videos, and experimental behavioral engagement: r = - 0.20, p = 0.03, N = 128 intervals for all 10 videos). Variability is measured as the standard deviation of the time estimates across viewers. Thus, people are not losing track of time, per se, but rather they are tracking time more similarly when they are engrossed in the story.

**Figure 3:**
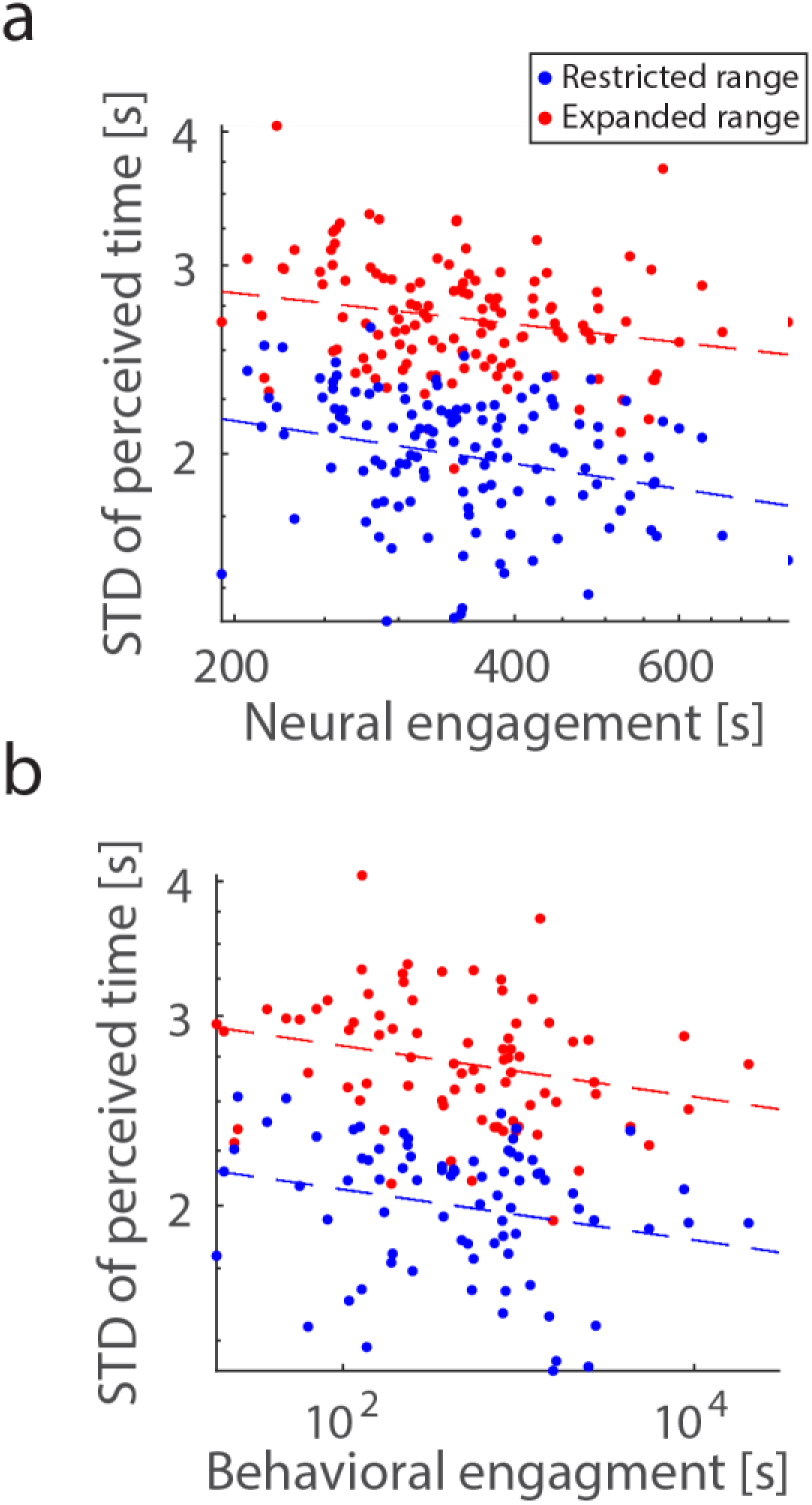
Engagement predicts the variability of time perception. Viewers estimated the duration of time intervals within each video. Each point represents a time interval in a video. **A:** The duration of intervals with high neural engagement were perceived less variably across viewers (r = - 0.27, p=2e-3, and r = - 0.23, p = 0.01, N = 129 intervals for all 10 videos, for restricted and expanded range, respectively). **B:** Higher real-world behavioral engagement also corresponded with a reduction in the variability of time perception (r = - 0.25, p=0.03, and r = - 0.25, p = 0.03, N = 78 intervals for the 5 real-world videos, for restricted and expanded range, respectively). All comparisons are made across two independent cohorts who had either a restricted range (blue, 8-16s) or expanded range (red, 4-20s) available for their time duration estimates. All time measures are displayed on a perceptual log-seconds scale.

To demonstrate the reproducibility of the findings, and because the distribution of time estimates was truncated in this initial experiment (see Supplementary Fig. S5), a second cohort was recruited. This cohort reported duration within an expanded range of 4s to 20s (“Expanded range”). The variability of time estimates across viewers again correlated with neural engagement (Figure 3A, Expanded range, r = - 0.23, p = 0.01, N = 129) and with behavioral engagement, measured in the real-world cohort (Figure 3B, Expanded range, r = - 0.25, p = 0.03, N = 78 intervals from the 5 real-world video). This weaker correlation was not resolved for experimental behavioral engagement (r = - 0.09, p = 0.3, N = 128 intervals from all 10 videos), possibly because the experimental data has a smaller range and is a noisier measure than the real-world assessment (compare Supplementary Fig. 1). Nevertheless, this independent cohort largely confirms our main and novel finding that engagement synchronizes neural activity across brains, thus resulting in a more uniform perception of time across people.

## Discussion

It is often said that we lose track of time when absorbed in an engaging narrative ^5,6^. Indeed, engagement can potentially either shorten or elongate our subjective perception of time ^7–9^. Engagement with a narrative may do both depending on its emotional valence ^10^. The precise neural processing that results in time’s distortion during naturalistic narrative stimuli is a matter of ongoing exploration ^14^. To determine the dependence of time perception on engagement, we first characterized engagement behaviorally in terms of how an audience is retained by narrative videos (Figure 1). To assess attentional engagement, we measured the inter-subject correlation (ISC) of stimulus-evoked EEG responses to the same videos (following ^15,16^). We found that ISC is predictive of the time that viewers were willing to spend with the videos (Figure 2). Surprisingly, neither behavioral nor attentional engagement correlated with the perceived duration of intervals of time during the videos. Instead, when the videos were more engaging, they were processed in the brain more uniformly (resulting in higher ISC), and the perception of time was more uniform across viewers (Figure 3). Thus, rather than losing track of time, viewers have a more consistent perception of time when highly engaged in the stimulus, at least for narrative videos on the scale of seconds.

We found that fluctuations in engagement behavior over time are predicted by the neural processing of the stimulus. The time-varying neural predictor, *γ*(*t*), modified the baseline level of engagement, *E*_0_, to track engagement behavior over time (Equation 2). In models that explore the survival of a population, the variable *γ* determines the “speed” at which the population ages, therefore modifying the expected life-span ^23^. It was therefore possible that while commitment to the stimulus remains constant, the subjective perception of time was variable. In this view, *γ*(*t*) would reflect the subjective perception that time “flows” differently depending on how immersed one is in the narrative ^17^. Alternatively, behavioral engagement could be interpreted in the context of “failure analysis” ^19^, where inverse risk, our definition of engagement behavior, quantifies the mean time to failure. In this case, inverse risk is the average time that it takes to lose a viewer. This idea is consistent with the interpretation of engagement as a time commitment, irrespective of the perception of time duration. The third experiment tested the two alternatives: whether engagement reflects time distortion or time commitment. We found that the perceived passage of time does not correlate with behavioral engagement nor its neural correlate. In contrast, both engagement measures correlated with the willingness to invest more time with the stimulus. Thus, our results are consistent with an interpretation of engagement as a value based decision, and not in terms of a perceptual warping of time. Indeed, time perception appears to have been less “warped” during moments of high engagement.

Despite extensive research on the neural underpinnings of time perception, no consensus exists on how humans estimate time intervals ^24^. While it is likely that different mechanisms support the perception of different time scales ^11^, there is also evidence for shared neural processes that implicate a diverse set of brain regions depending on task specifics ^18^. The prominent models, such as the pacemaker-accumulator models ^25^, and the climbing neural activation model ^24^, posit that there is some input that is integrated across time, and that this integrated signal outputs an estimate of duration. Previous fMRI studies have explored the dependence of such signals on attention ^26^, emotional content ^27^, and salience ^28^. It is likely that all of these features altered time perception during our video stimuli. We found that more similar neural processing across viewers led to a more uniform report of time’s passage. This suggests that stimulus processing may ultimately provide (or at least modulate) the input to the time integration circuit, because more uniform stimulus processing (input) led to more uniform time estimates (output) across viewers. It is relatively unique to measure time perception during dynamic natural stimuli. Recently, Lositsky et al. (2016) measured time perception during auditory narratives and found that changes in stimulus induced brain activity are predictive of retrospective time estimates on the order of minutes. Our results concern prospective time estimates for intervals on the order of seconds. Despite these differences, both studies support the notion that for narrative stimuli, the subjective passage of time is driven by stimulus processing, rather than an internal stimulus-independent “clock” ^29^.

It is well established that time perception is affected by attention ^13^. We therefore leveraged EEG responses evoked by the stimuli to gauge attentional engagement using ISC. The brain activity most predictive of engagement was the second component of the EEG activity correlated across viewers (see Figure 3A for scalp topography). With a factor of *γ*_2_ = 1.5, this component explains the fluctuation in time commitment from a baseline level of approximately 200s to the average engagement level of approximately 300s. Interestingly, among the three strongest components of the ISC, this component is also the best at discriminating between attentional states ^15^. This component is also uniquely evoked during narratives with sound, as opposed to those with only visuals, and it may therefore be representative of auditory processing ^20^, or potentially how strongly the stimulus’ auditory narrative captures attentional resources.

The concept of “engagement” is used in a variety of contexts: a client engages a law firm, a couple is engaged to be married, or a student is engaged in the classroom. In all of these scenarios there is a commitment of either time or financial resources. Similarly, in this paper, we define engagement as the commitment of a scarce resource. Unlike self-report assessments ^5,17^, this definition is quantifiable in strict numerical terms and can be applied even when subjective reports are impossible ^30^. Whether the resources engaged are time or attention, a value-based decision is consistently assessed in which the value gained from consuming the narrative is compared to that of possible alternatives ^31^. Here, both our neural and behavioral measures of engagement predict that viewers can commit time periods that range from seconds to hours (see units on Figure 2B). In the third experiment, this commitment was literally represented by the decision to spend time with the videos rather than to earn money. This is because the subjects, recruited on MTurk, were forfeiting potential income from the completion of the tasks for which they are paid by continuing to watch the videos. The correlation of both engagement measures with this decision is a direct confirmation that engagement is a value-based decision (Figure 1C and 2C).

To evaluate engagement behavior, an experimental technique was developed that takes a fraction of the time necessary to acquire comparable real-world data. Our definition of engagement, in terms of devoting a scarce resource, and potentially this technique, could be translated to other media that engage people such as books, music, virtual reality, gaming, or to stimulating activities such as painting or playing sports. Time does not necessarily have to be the resource that is sacrificed. By making a commitment, viewers may be foregoing monetary compensation or social rewards to gain access to entertainment. The worth of engagement may be computed using a currency that can be traded for these and other scarce resources. We predict that if viewers are similarly entrained by the stimulus (or activity), thus eliciting a high level of ISC, they will be immune to extrinsic costs such as the time or money that they are sacrificing for the current moment’s enjoyment. Their perception of time, one of the many valuable resources that they are sacrificing, will thus be driven by the stimulus, and consistently so across viewers.

## Methods

### 1. Stimuli

Stimuli were chosen on the basis of their highly emotive content and the availability of online viewership data. We used these same stimuli in a previous EEG study on incidental memory ^20^. They consisted of 10 different videos (5 from the New York Times' Modern Love episodes: “Broken Heart Doctor”, “Don't Let it Snow”, “Falling in Love at 71”, “Lost and Found”, and “The Matchmaker”, and 5 from StoryCorps' animated shorts: “Eyes on the Stars”, “John and Joe”, “Marking the Distance”, “Sundays at Rocco’s” (depicted in Figure 1A), and “To R.P. Salazar with Love”). Stimuli were 161 ± 44 s in duration (mean and standard deviation across stimuli) with a total duration of 26 min 48 s.

### 2. Behavioral Engagement Data Collection

Real-world viewing behavior was measured using the pool of viewers who found the five StoryCorps videos organically on YouTube, via the StoryCorps website (storycorps.org/animation), or another linked website. Anonymous YouTube Analytics data were provided as aggregated viewership survival data (Figure 1A, Supplementary Fig. 1) by StoryCorps with permission for analysis and publication. The viewership data captured the behavior of viewers, totaling 2,528,897 across all five videos, amassed since the videos were made available online until the time of data retrieval (2.6 +/- 0.9 years).

Behavioral engagement was measured experimentally on an independent set of 1,000 subjects collected on Amazon’s Mechanical Turk (MTurk) platform (requester.mturk.com). Participants with IP addresses located in the USA had access to all 10 stimuli in a randomized order for a duration of 15 minutes and were informed that the total duration of all videos exceeded the time allotted. Watching the videos was optional (required a mouse click) and it is therefore possible that a fraction of these participants did not watch any of the videos. It took approximately one hour for this data to be collected for all 1000 subjects. This collection method was selected after pilot testing in three smaller cohorts of 100 subjects each. In the first pilot, participants had the option of immediate remuneration and therefore spent little time with the videos. In the second pilot, participants were not informed that they would be paid. In the third pilot, subjects were informed of their impending payment. The results of this pilot agreed best with the real-world data, and this format was therefore used for the final data collection. None of the pilot data was included in the final analysis reported here.

The YouTube Analytics API was used to extract the fraction of viewers (number of current viewers / total number of initial viewers) who watched each video interval. The resulting curve can be considered the viewership survival, S(t), although it is not a strict survival metric because YouTube allows users to rewind and skip sections and therefore S(t) sometimes increases. Note that the Analytics API divides each video into 100 sampling intervals, regardless of video duration, and was resampled to correspond with the absolute time elapsed, as described below.

### 3. Risk of viewership loss

At each time interval, a decision is made regarding whether the current activity provides more reward than another activity. In aggregate, over the population of viewers, this is reflected in the survival function, *S*(*t*), the ratio of people that are still watching, or have survived, until time*t*. A typical example of this from the real-world data is shown in Figure 1A. The risk, or hazard, of losing viewers, *λ*(*t*), can be estimated from *S*(*t*)and is conventionally defined as the relative change of the survival in a time interval *Δt* ^23^:

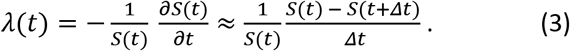

In a realistic scenario, stimuli are selected, engaged with, and finally abandoned when audience members determines that their limited temporal resources are better spent elsewhere. Figure 2A shows several real-world examples of how the number of surviving viewers, *S*(*t*),decays over time. The hazard curve, *λ*(*t*), derived from the survival function (Supplementary Fig. S1) presents a typical “bathtub” curve with high risk of viewership loss at the beginning and ending and low risk during the intervening time. In failure analysis inverse risk represents the mean time to failure ^19^, and for constant risk, the survival curve is exponentially decaying with the inverse risk as its time constant.

Risk of losing the audience, *λ*(*t*), was computed according to Equation (3) on the 100 samples in the raw data. For visualization purposes, Figure 1A and Supplementary Fig. S1 show survival, S(t), and risk, *λ*(*t*), as a function of absolute time (scaled to seconds). For the purpose of comparing experimental and real-world data, *λ*(*t*) was resampled to a time scale of ∆t =12s. Engagement, E(t), derived from *λ*(*t*)in Equation (1), is plotted at this scale (Supplementary Fig. S1). This time scale was motivated by the comparison with the neural data. To calculate the average number of viewers lost in a time interval of ∆t =12s, the derivative of S(t) was scaled by the initial number of viewers and then multiplied by 12 seconds.

Time intervals with negative engagement (due to rewinding and skipping on YouTube) are ignored because the logarithm of engagement is used in all analyses. Therefore, at a sampling interval of ∆t =12s, 1 interval is excluded from the experimentally acquired data, and 3 intervals are excluded from the real-world data. This exclusion effects the comparison between the two behavioral engagement measures (Figure 1B, a reduction from N=81 intervals to N=77), the comparison with the decision to continue watching (Figure 1C, a reduction from N=122 to N=121), the comparison with neural engagement (Figure 2B, Results Section 2, for the experimental cohort a reduction from N=129 to N=128, and for the real-world cohort a reduction from N=81 to N=78), and the comparison with the time estimates (Figure 3B, reductions the same as those for neural engagement).

### 4. Perceived Time Data Collection

Perceived time was measured on two additional MTurk cohorts. Each participant watched all 10 stimuli and a pseudo-randomly selected 12 second time interval was denoted by the appearance of a red hourglass in the corner of the video. After the interval had expired, the video was paused, and participants were asked to estimate the duration for which the hourglass had appeared (perceived time). The longest video clips had 19 intervals. To ensure that each of these intervals had at least 20 estimates, 380 subjects were recruited. Across all videos, there were N=129 time intervals, of these 129, N=81 corresponded to intervals in the videos for which real-world behavioral engagement was assessed. In this first cohort, subjects reported perceived duration on a visual analog scale with values ranging from 8 to 16 seconds. A histogram of reported time estimates shows a large number of responses at the boundaries of this range (see Supplementary Fig. S5, restricted range), suggesting that inputs may have been restricted. A second cohort was therefore recruited and given the option to input estimates ranging from 4 to 20 seconds (expanded range). To ensure a robust estimate of the standard deviation of perceived time across subjects, this cohort had 720 participants. After each interval had transpired, participants were also asked a comprehension question related to the content of the story in that interval (same questions as in ^20^; mean accuracy level 84.0% +/- 37.7% across questions). After the response was recorded, participants were given the option to finish the video. When the interval that was just watched was the last interval in the video, the option to finish the video was not given. As this occurred in 7 cases, this resulted in a reduction from N=129 intervals to N=122 intervals. Prior to the presentation of the 10 experimental stimuli, participants were briefly acquainted with the task on four non-experimental videos. For the first video, subjects are told ahead of time that the interval is 12 seconds in duration; for the next three, interval durations are selected at random to be either 8, 12 or 16 seconds. Without prior knowledge of the duration, subjects are asked to estimate it and are subsequently informed of the correct duration. All data collection was approved by the Institutional Review Board of the City University of New York.

### 5. Electroencephalographic Data Collection and Preprocessing

Electroencephalographic (EEG) data were collected for a previous study from a cohort of 20 individuals who watched the same audiovisual stimuli (AV condition, in Cohen & Parra, 2016). For more details regarding participants, EEG data collection, and preprocessing see Cohen & Parra, 2016.

### 6. Inter-Subject Correlation

Inter-subject correlation (ISC) is evaluated in the correlated components of the EEG ^16^. The goal of correlated component analysis in this case is to find linear combinations of electrodes (one could think of them as virtual sensors or “sources” in the brain) that are maximally correlated between subjects. The ISC calculation implemented here is the same as what has been published previously ^15,20^. Following previous research, the top three components of the EEG are extracted and capture maximally correlated responses between subjects ^16^. To determine the correspondence between behavioral engagement and ISC, ISC was resolved in time, using the same time intervals over which behavioral engagement was measured (see next section). Time resolved ISC in the *i*-th correlated component is here denoted as*x*_*i*_(*t*).

### 7. Comparisons between behavioral and neural engagement

A proportional hazard model was used (Cox, 1972) to relate engagement behavior and ISC. This resulted in a regression of engagement, *E*(*t*), with a time dependent covariate, *γ*(*t*), and a constant baseline engagement, *E*_0_ (Equation 2 in Results). Following the traditional form of the proportional hazard model ^19,32^, the time dependent covariate, *log γ*(*t*), equals the weighted sum of the predictor variables:

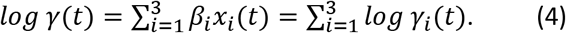

The contribution of ISC in each component, *x*_*i*_(*t*), to the total engagement can thereby be assessed from its corresponding *γ*_*i*_(*t*)value. Optimal model parameters *β*_*i*_ and *E*_0_ were found as the best linear fit for the log-engagement data collected from the experimental group. This fit was performed separately for different time intervals (∆t). Engagement, *E*(*t*), was resampled from the 100 samples in the raw data into various time resolutions (∆t = 1s, 2s, 3s, … and up to 30s) and time-resolved ISC *x*_*i*_(*t*) was also calculated on the matching time intervals. Goodness of fit, R, is shown in Supplementary Fig. S3 as a function of ∆t. For consistency, all subsequent analyses were performed at 12s resolution, which provided the best fit (see Results).

Final parameter estimates were assessed by training on the experimentally measured engagement data and testing on the real-world data without further parameter adaptation. This captures the generalization performance from trained to unseen data, as well as generalization from the experimental procedure to the real-world behavioral engagement data. To capture the generalization between different stimuli, in an additional analysis, training of the model parameters, *β*_*i*_, and *E*_0_, was limited to only those videos that were not in the testing set, i.e. training was performed on the experimental data from the 5 New York Times videos and the real-world data from the 5 StoryCorps videos was used for testing.

### 8. Relationship between time perception and engagement

Correlation was assessed between both engagement measures (behavioral engagement: *log*(*E*(*t*)) and neural engagement: using the optimal fit for *β*_*i*_ at ∆t=12s) and the perceived time of each interval, collected from two independent cohorts of subjects. As expected ^13^, mean time perception was longer for later intervals in the videos (Supplementary Fig. S4). This drift in mean time perception with wait time does not affect the standard deviation of time estimates because the standard deviation does not depend on the mean (Figure 3). Linear regressions were used to relate the standard deviation of time estimates to both neural and behavioral measures of engagement.

### 9. Statistics

The significance of reported correlation values, r, were computed using conventional parametric statistics which assume independent samples. To rule out that temporal correlations between samples bias these results, p-values for all analyses were also estimated using nonparametric bootstrapping. Specifically, significance levels were computed using phase shuffled data (following ^33^), which preserve the correlation structure in time but alters the time course of a temporal sequence. Therefore, N=10^6^ phase shuffled surrogates were produced and correlation coefficient computed for each surrogate. Bootstrap p-values are calculated as the fraction of shuffles with correlation values more extreme than those obtained with the original time sequences. All bootstrap p-values were comparable to the values reported in the main text using parametric statistics (except for the comparison between real-world behavioral engagement and the variance of time estimates, Figure 3C where p > 0.2). In the analysis of the viewer's decision to continue to watch the videos (“Fraction continued video” in Figure 1C, and 2C), data from the two time estimation cohorts (Restricted and Expanded) was combined since the decision to continue watching did not vary across the cohorts. All analysis of time and engagement were performed on a log-seconds scale (except for the computation of the mean and standard deviation of the time estimates, Figure 3). All statistical tests were performed in MATLAB (MathWorks, Natick, MA, USA).

## Acknowledgements

We would like to acknowledge StoryCorps for providing us with their YouTube analytics data and their animated interviews as video stimuli. These interviews were recorded by StoryCorps and are provided courtesy of StoryCorps, a national not-for-profit corporation dedicated to preserve and share humanity's stories in order to build connections between people and create a more just and compassionate world. www.storycorps.org.

